# A deep network-based model of hippocampal memory functions under normal and Alzheimer’s disease conditions

**DOI:** 10.1101/2021.01.31.429076

**Authors:** Tamizharasan Kanagamani, V. Srinivasa Chakravarthy, Balaraman Ravindran

## Abstract

We present a deep network-based model of the associative memory functions of the hippocampus. The proposed network architecture has two key modules: 1) an autoencoder module which represents the forward and backward projections of the cortico-hippocampal projections and 2) a module that computes familiarity of the stimulus and implements hill-climbing over the familiarity which represents the dynamics of the loops within the hippocampus. The proposed network is used in two simulation studies. In the first part of the study, the network is used to simulate image pattern completion by autoassociation under normal conditions. In the second part of the study, the proposed network is extended to a heteroassociative memory and is used to simulate picture naming task in normal and Alzheimer’s disease (AD) conditions. The network is trained on pictures and names of digits from 0 – 9. The encoder layer of the network is partly damaged to simulate AD conditions. As in case of AD patients, under moderate damage condition, the network recalls superordinate words (“odd” instead of “nine”). Under severe damage conditions, the network shows a null response (“I don’t know”). Neurobiological plausibility of the model is extensively discussed.

## 1. Introduction

There is a long line of studies that implicate the role of the hippocampus in declarative memory functions (De Almeida, Idiart, and Lisman 2007; Steinvorth, Levine, and Corkin 2005; Brenda Milner, Corkin, and Teuber 1968). Damage to the hippocampal region is seen during the course of Alzheimer’s disease and the normal course of aging (Golomb et al. 1993). Initial understanding of the medial temporal lobe and its substructures, especially the hippocampus, in memory functions came from the studies on the famous patient HM, whose name came to be indelibly linked to hippocampal memory studies (De Almeida, Idiart, and Lisman 2007; Penfield and Milner 1958a; B. Milner 1966; MILNER 1970). Scoville et al. (1957) reported severe anterograde amnesia and moderate (less severe) retrograde amnesia in HM. However, memories from long before the point of injury were intact, which showed that the hippocampus is only a temporary store of memory and that memories are subsequently transferred out of the hippocampus to long-term stores. This process is known as consolidation (McGaugh 2000; Diekelmann and Born 2010; Born and Wilhelm 2012; Marshall and Born 2007). HM showed normal behavior in the immediate recall but had performance impairment in delayed recall tasks (Baddeley and Warrington 1970). Although HM failed to learn new episodic information, he had no problem learning motor skills such as visual-motor tracking tasks (Corkin 1968). Further confirmation came from animal studies that demonstrated that removal of the hippocampus leads to impairment in the formation of new memories (Terzian and Ore 1955; Penfield and Milner 1958a). Yonelinas et al. (2004) and Fortin et al. (2004) showed that the damage to the hippocampus leads to the impairment in associative recall (A. P. Yonelinas et al. 2004; Fortin, Wright, and Eichenbaum 2004).

In order to serve its function as a memory unit, the hippocampus must have access to the raw material for memory, which is sensory information. A quick review of the anatomy of the hippocampus and its place vis a vis the cortex provides useful insights into the mechanisms of its memory functions. As a subcortical circuit, the hippocampus receives widespread projections from cortical areas in the temporal, parietal, and frontal lobes (E 1999; R. Insausti et al. 2017). A majority of hippocampal afferents from the posterior brain come from higher-order sensory and association cortices, areas that are capable of generating abstract representations of sensory information (Bowman and Zeithamova 2018). Thus, sensory information spread out over large cortical areas is projected, first to parahippocampal and perirhinal cortices, and then to the entorhinal cortex (EC), which is the gateway to the hippocampus (Burwell and Amaral 1998). Neurons of EC then project to small neural clusters of the hippocampal circuit, making it possible to create compact representations of elaborate cortical states (Menno P Witter 1993). There are also back projections from the hippocampus to the same cortical targets, thereby creating the possibility of reconstructing the cortical state based on the more compact hippocampal representations (Rosene and Van Hoesen 1977; Catenoix et al. 2011). Such a reconstruction is thought to be the basis of memory *recall* (Linde-Domingo et al. 2019; Chrobak, Lrincz, and Buzsáki 2000).

The projection pattern from cortical areas to the hippocampus suggests that one of the prime features of the cortical state represented by the hippocampus is *pattern separation* (Yassa and Stark 2011). To illustrate the concept of pattern separation, let us consider the problem of representing a cricket ball vs. a tomato. A cricket ball is round, red, and hard, while tomato is approximately round, red, and soft. Thus, the cortical representations of the two objects are likely to have a large overlap. But since the objects these feature combinations point to are quite distinct, it is desirable that the representations generated by the hippocampus are also adequately distinct, thereby achieving pattern separation.

There is another aspect of hippocampal memory function known as *pattern completion* (E. T. Rolls 2013; Mizumori et al. 1989). Let us illustrate this concept using the same objects: a cricket ball and tomato. When we identify a cricket ball from a picture (red + round) even without touching it (hard), we are mentally supplying the missing feature of hardness. Similarly, a tomato can be identified visually without tactile exploration. These are examples of pattern completion that involves filling in missing features based on sensed features.

In order to understand how the hippocampus supports pattern separation and pattern completion, one must consider the anatomy of the hippocampus circuit. The hippocampal formation connects several neural fields like the Dentate gyrus (DG), CA3, CA1, and subiculum (C. Schultz and Engelhardt 2014). Nearly all the neural fields in the hippocampus receive projections from the superficial layers of the Entorhinal Cortex (EC) (Brun et al. 2008; Hargreaves et al. 2005; Gloveli, Schmitz, and Heinemann 1998). The EC, in turn, receives projections from widespread cortical areas in the prefrontal, parietal, and temporal lobes via several intermediary regions and also sends back projections to the same areas (Kondo and Zaborszky 2016; Agster and Burwell 2009). EC’s afferent connections are formed using one trisynaptic pathway and two monosynaptic pathways (Yeckel and Berger 1990; Charpak, Paré, and Llinás 1995). The trisynaptic pathway consists of the perforant pathway between the second layer of EC (EC II) to DG (M. P. Witter, Van Hoesen, and Amaral 1989), the mossy fibers between DG and CA3 (Claiborne, Amaral, and Cowan 1986), and Schaffer collaterals between CA3 to CA1 (Kajiwara et al. 2008). The monosynaptic pathways are formed between the second layer of EC (EC II) to CA3 (Gloveli, Schmitz, and Heinemann 1998; Empson and Heinemann 1995) and the third layer of EC (EC III) to CA1 via perforant pathways (M. P. Witter, Van Hoesen, and Amaral 1989). CA3 has more recurrent connections compared to the other hippocampal regions (D. G. Amaral and Witter 1989), a feature that qualifies it for a crucial role in pattern completion and memory storage. The fifth layer of EC (EC V) receives the afferent projections from CA1 directly and indirectly via the subiculum (Canto et al. 2012; K. C. O’Reilly et al. 2013). It is this fifth layer of EC that sends back projections to widespread cortical targets (Ricardo Insausti, Herrero, and Witter 1997).

A majority of hippocampal memory models involve implementations of pattern separation and pattern completion, distinguishing themselves in terms of anatomical details incorporated in the model or the specific memory tasks that they set out to explain (Marr 1971; R. C. O’Reilly and McClelland 1994; E. T. Rolls and Treves 1994; McNaughton, Connectionist, and 1990 2020; E. Rolls and Treves 2012). Gluck and Myers (1993) exploit the cortico-hippocampal projection pattern (Gluck and Myers 1993), which they model as an autoencoder (G. E. Hinton 2006). The nature of the autoencoder training ensures that, in this model, the hippocampus achieves pattern separation. The high-dimensional cortical state is the input to the autoencoder, while the hippocampus is the low-dimensional hidden layer. Thus, the cortico-hippocampal projections form the encoder, while the back projections representing the decoder are responsible for memory recall.

The pattern completion aspect of hippocampal memory function was highlighted by one of the earliest and most influential models of the hippocampus proposed by David Marr (1991) (Marr 1971). Marr visualized a memory as a pattern distributed over a large number of neocortical neurons. Since the neocortical neurons have reentrant connections, it is possible to store patterns by association. Activation of some neurons that represent a partial set of features can cause activation of neurons representing the remaining features, thereby achieving pattern completion. Mathematical associative memory models that exhibit pattern completion often involve networks with high recurrent connectivity and attractor dynamics (John J. Hopfield 2018; Amit 1990; J J Hopfield 1984). The high recurrent connectivity (4%) among the CA3 pyramidal neurons had inspired a long modeling tradition that treats CA3 as an associative memory (De Almeida, Idiart, and Lisman 2007; David G. Amaral, Ishizuka, and Claiborne 1990). Treves and Rolls (1994) have taken the associative memory view of CA3 and presented storage capacity calculations (Treves and Rolls 1994). Wu et al. (1996) described the effect of noise of pattern storage in an associative memory model of CA3 (Wu, Baxter, and Levy 1996). This has evolved a computational perspective that posits CA3 at the heart of pattern completion functions of the hippocampus.

There are other modeling approaches that describe pattern completion mechanisms of the hippocampus without specifically describing CA3 as an associative memory. The models of O’Reilly and colleagues describe the loop of connections from the superficial layers of EC, to DG to CA3 to CA1 back to deep layers of EC (Norman and Reilly 2002; R. C. O’Reilly and Rudy 2001; R. C. O’Reilly and McClelland 1994). Hasselmo and Wyble present a model of hippocampal attractor dynamics that explains the disruptive effects of scopolamine on memory storage (Hasselmo and Wyble 1997; Hasselmo, Wyble, and Wallenstein 1997). Thus, there is a spectrum of models that describe pattern completion functions of the hippocampus either by placing the burden of storage exclusively on CA3 and its recurrent connectivity or relying on the general internal loops of the hippocampus to supply the necessary attractor dynamics.

The aforementioned review of computational models of hippocampal memory functions shows a common structure underlying a majority of the models embodying two crucial features: 1) They impose some form of autoencoder structure, with feedforward/feedback projections, on the cortico-hippocampal network, thereby achieving pattern separation and a compact representation of the cortical state. 2) They use the attractor dynamics arising, either solely within CA3 or, more broadly, in the hippocampal loops to achieve pattern completion. Instead of addressing the sensitive task of having to pick the best among the above models, we propose to construct a model with the above features but cast in the framework of deep networks so as to exploit the special advantages offered by deep networks.

Although often criticized for possessing inadequate biological plausibility, in recent years, deep networks have enjoyed surprising success in modeling the activities of visual, auditory, and somatosensory cortical hierarchies (R. C. O’Reilly and McClelland 1994; Kanitscheider and Fiete 2017; Norman and Reilly 2002; R. C. O’Reilly and Rudy 2001). Some progress has been made in the use of deep networks for modeling hippocampal spatial navigation functions but not its memory functions (Kanitscheider and Fiete 2017).

In this paper, we present a deep network-based model of hippocampal memory functions. The network has a deep autoencoder structure with the inner-most layer, the Central Layer (CL), representing the hippocampus. Furthermore, attractor dynamics is imposed on the state of CL by assuming that the state of CL constantly seeks to find the local maximum of a *familiarity* function, where the familiarity refers to the confidence at which an object is remembered (Skinner and Fernandes 2007). The proposed network exhibits more accurate recall performance than one without the attractor dynamics over the familiarity function. Going beyond the basic model, we implement the picture-naming task by introducing two pipelines in the network architecture – one for the image and another for text. We apply the resulting “picture naming model” to simulate the performance of Alzheimer’s patients. When the hidden layer neurons are progressively destroyed, in order to simulate hippocampal damage in Alzheimer’s, the model’s recall performance showed a strong resemblance to the performance of the patients on the same task.

## 2. Methods and Results

### 2.1 Auto-Associative Memory Model

The model of auto-associative memory is explained using a modified convolutional autoencoder, in which the Central Layer is associated with attractor dynamics. We call such an architecture an Attractor-based Convolutional Autoencoder (ACA). The attractor dynamics arises out of performing hill-climbing over a cost function, which in the present case is the familiarity function. The performance of the proposed model is compared with a standard convolutional autoencoder and a recurrent convolutional autoencoder. All the three architectures are compared on image pattern completion tasks.

#### 2.1.1 Standard Convolutional Autoencoder (SCA)

The standard convolutional autoencoder network (Figure 1) takes an image as input and maps it onto itself, or often to a noise-free version of itself. In the present case, the encoder comprises two convolution layers with max-pooling followed by two fully connected layers. The decoder comprises two fully connected layers followed by two deconvolution layers, thereby producing the output of the same size as the input.

**Figure 1.**
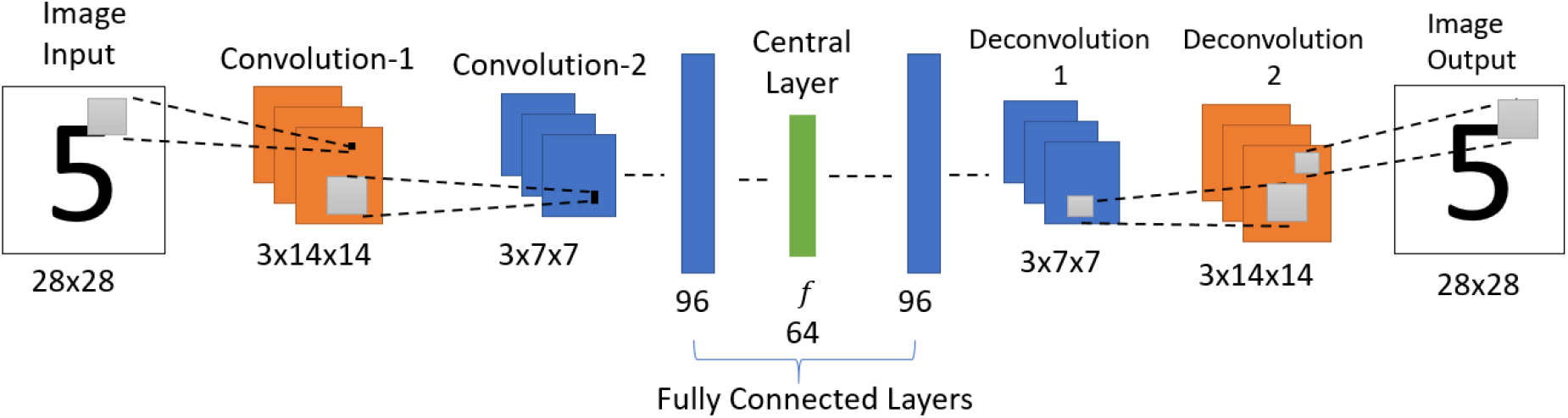
Architecture of Standard Convolutional Autoencoder.

#### 2.1.2 Image Encoder

The encoder uses the input image of dimension 28×28. It comprises two convolution layers, with each convolution layer extracting three feature maps over a receptive field of size 3×3. Each convolution layer is followed by a max-pooling layer (Scherer, Müller, and Behnke 2010) of stride 2 generating feature maps, each of size 3×14×14 and 3×7×7, respectively. The output of the second convolution layer is flattened (3×7×7 to 147) and connected to a fully connected layer with 96 neurons. This, in turn, is connected to the Central Layer with 64 neurons. Here all the layers use the leaky-ReLU activation function (Maas, Hannun, and Ng 2013), but the Central Layer uses the sigmoid activation function.

#### 2.1.3 Image Decoder

The image decoder generates an image using the features from the Central Layer. Here, the final layer of the encoder is connected to a fully connected layer with 96 neurons. This, in turn, is connected to a fully connected layer with 147 neurons, which is then reshaped to 3×7×7. This reshaped data is fed to the first deconvolutional layer, which has three filters of size 3×3 with stride 2 and produces an output of size 3×14×14. Then the first deconvolutional output is fed to the second deconvolutional layer (3 filters of size 3×3 with stride 2) to produce the image output of size 28×28. Here the output layer uses a sigmoid activation function, and all the other layers use leaky-ReLU activation function.

#### 2.1.4 Recurrent Convolutional Autoencoder (RCA)

This model uses essentially the same architecture as the one in the standard convolutional autoencoder network above but with a difference: instead of decoding the input in one-step, the output of the network at the current iteration is used as input in the next iteration (Figure 2). Thereby forming a loop, the network acts as an attractor, and the stable output obtained after several iterations is considered as the final retrieved pattern.

**Figure 2.**
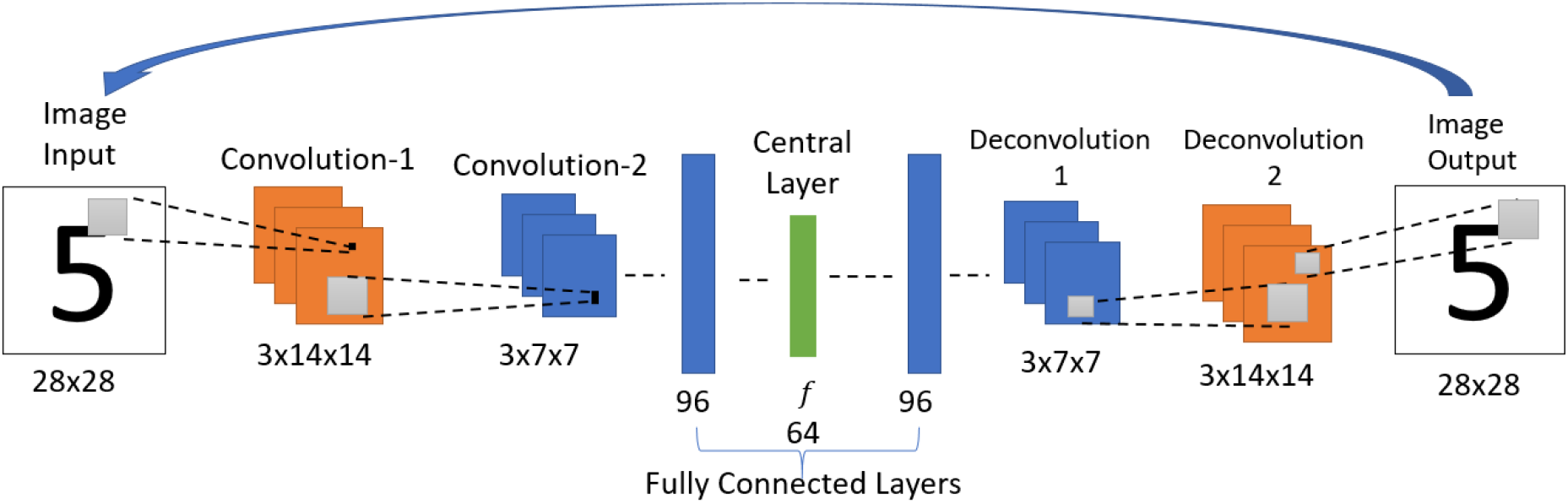
Architecture of Recurrent Convolutional Autoencoder

#### 2.1.5 Attractor-based Convolutional Autoencoder (ACA)

This model also uses the same network architecture as used in the simple convolutional autoencoder above, but with an important modification (Figure 3). Here the architecture of the encoder and the decoder are the same as used in the simple convolutional autoencoder network. This model uses the notion of familiarity, and each input to the network is mapped to a scalar value that represents familiarity. Here we hypothesize that the EC is estimating the familiarity value, and the hippocampus uses it to recall the memory. In the network, the “familiarity unit” is implemented by a single sigmoidal neuron, which receives the inputs from the central layer (CL) with 64 neurons (attributed to EC) via a sigmoidal layer with 32 neurons. This unit outputs a scalar value V representing the familiarity level of the input. This single node predicts the familiarity (correctness) value *C* (equation (1)).

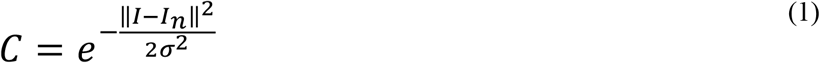

where *I_n_* denotes the noisy version of an image. *I* represents the perfect/noiseless Image. Here σ is set to 15. Whereas the value *C* for a noiseless image is 1, for a noisy image, the value ranges between 0 and 1. For each training image, the corresponding value is calculated using equation (1). After training, the familiarity value predicted by the network is used to reach the nearest best feature with the maximum familiarity value.

**Figure 3.**
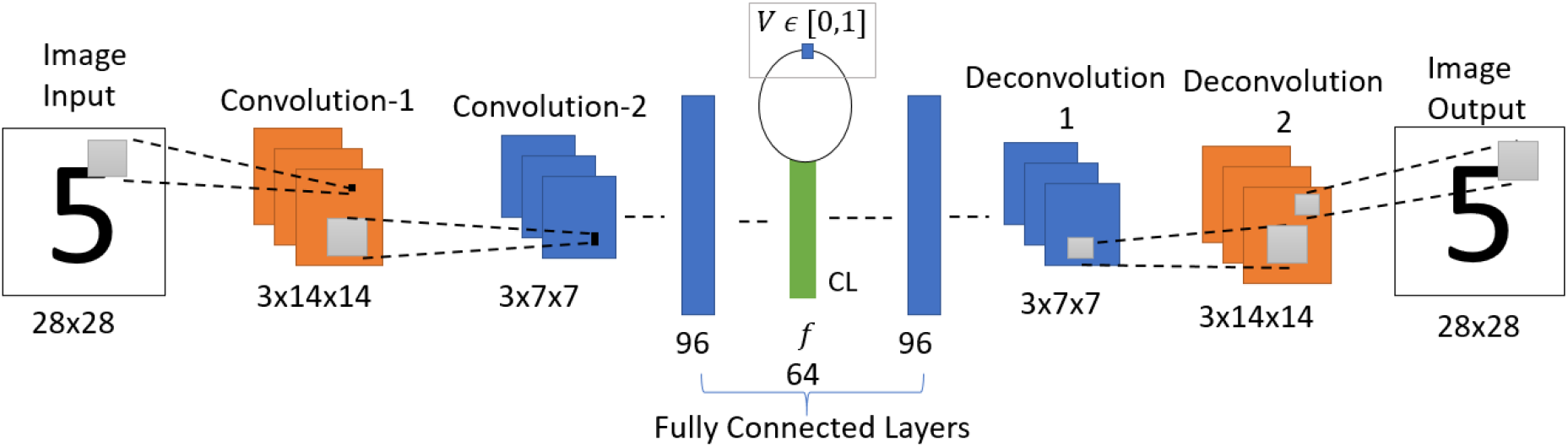
Architecture of Value-based Convolutional Autoencoder Network. CL-Central Layer

#### 2.1.6 Dataset

The image dataset is generated using the images of printed numerals 0-9 (Figure 4) with size 28×28. The dataset consists of images with various noise levels. The noisy images are generated using equation (2).

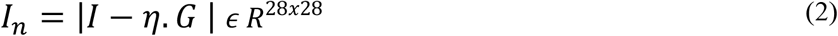

**Figure 4.**
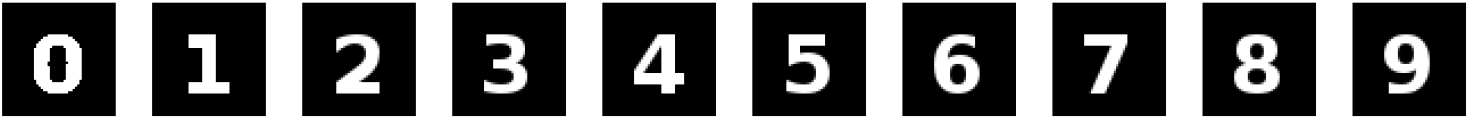
Digit Images without noise

Here *I*, and *I_n_* denotes the noiseless Image, noisy Image, respectively. *G* represents the noise matrix, whose individual element is given as *G_ij_* = *U*(0,1). Where *U*(0,1) is a uniform random variable with values ranging between 0 and 1. *η* is a scalar that specifies the noise percentage, which ranges between 0 and 1. The modulus operator is used to keep the image pixel values between 0 and 1. The noisy sample images are shown in Figure 5. The training dataset contains 1,00,000 images, the validation dataset contains 20,000, and test dataset contains 20,000 images. The images are categorized into ten classes (0-9) depending on the source image it is generated.

**Figure 5:**
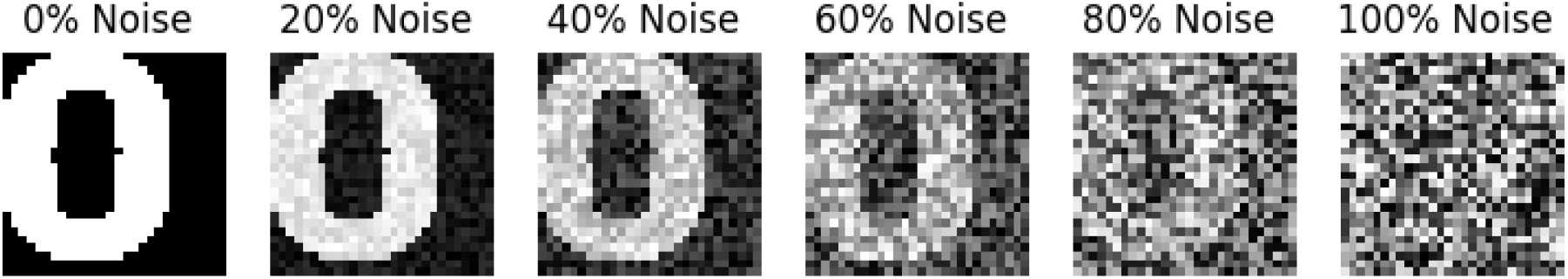
Image of Zero at different noise levels.

#### 2.1.7 Training

##### 2.1.7.1 Standard Convolutional Autoencoder

Unlike the other denoising autoencoders, the standard convolutional autoencoder is trained to produce the same noisy input as output. The standard autoencoder network is trained to minimize the cost function 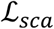 (a combination of multiple cost functions) as given below.

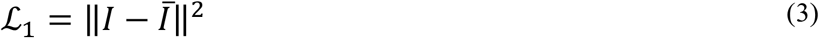

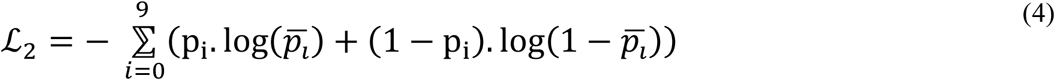

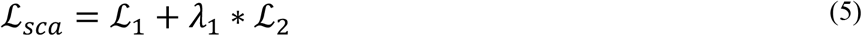

Here,

*I* – input image
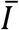 - predicted Image
p_i_ - actual probability of input image being in i^th^ class
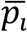 - predicted probability of being in i^th^ class
*λ*_1_ – trade-off parameter
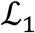 denotes the image reconstruction error. 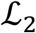 denotes classification error (cross-entropy). SoftMax layer with ten neurons is used for classification over the Central Layer. The classification layer is used to make the encoded features separable for the image inputs of different numbers. The network parameters are updated using Adam optimizer (Kingma and Ba 2015).

##### 2.1.7.2 Recurrent Convolutional Autoencoder (RCA)

As the standard convolutional autoencoder itself is merely used iteratively, there is no separate training employed in this case.

##### 2.1.7.3 Attractor-based Convolutional Autoencoder (ACA)

The Attractor-based convolutional autoencoder is trained similarly to the standard convolutional autoencoder along with an additional cost function for the familiarity value prediction. Here the attractor-based convolutional autoencoder is trained to minimize the cost function 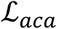 (Equation (7)).

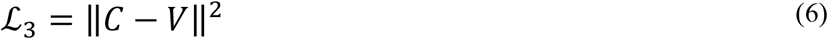

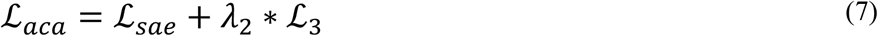

Here,

*C* – desired familiarity value
*V* – predicted familiarity value
*λ*_2_ – tradeoff parameter
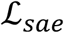 - the cost function used in the standard convolutional autoencoder.
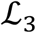 denotes the familiarity value prediction error. Here also the network is trained using Adam optimizer.

In this network, for a given input image, the output is retrieved after modifying the encoded feature using the familiarity value. The feature is modified using the predicted familiarity value to attain the maximum familiarity value by the hill-climbing technique. Here Go-Explore-NoGo paradigm is used to implement the stochastic hill-climbing behavior.

#### 2.1.8 Go-Explore-NoGo (GEN)

Similar to Simulated Annealing (Kirkpatrick, Gelatt, and Vecchi 1983), the GEN algorithm allows us to perform hill-climbing over a cost function without explicitly calculating gradients. Although originally derived in the context of modeling the basal ganglia, it can be used as a general optimizing algorithm. The Go-Explore-NoGo (GEN) policy (Srinivasa Chakravarthy and Balasubramani 2018), consists of 3 regimes: Go, Explore, and NoGo. A slightly modified version of GEN is used in this model. The Go regime decides that the previous action must be repeated. The NoGo regime forbids from taking any action. (There is another variation of the NoGo regime wherein the action taken is opposite of the corresponding action in the Go regime (Srinivasa Chakravarthy and Balasubramani 2018)). The Explore regime allows choice of a random action over the available action space. For a given input image, the feature *f* from the Central Layer and the corresponding familiarity value *V* is estimated. The network aims to identify the nearest feature with a maximum familiarity value of 1. It is achieved using the following algorithm based on the GEN policy (Srinivasa Chakravarthy and Balasubramani 2018).

Let

*f* - be the current feature vector for the given input image
*V* - familiarity value for the feature vector *f*
*ϵ* - threshold value
ϕ - is a 64-dimensional random vector, where each element ϕ_i_ is given as.

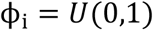 Where *U*(0,1) is a standard uniform distribution variable with values between 0 and 1. Initialize Δ*f*(0) = 0
*f*(*t* + 1) = *f*(*t*) + Δ*f*(*t*)
*δ*(*t*) = *V*(*t* + 1) – *V*(*t*)
*f* is updated by the following equations:

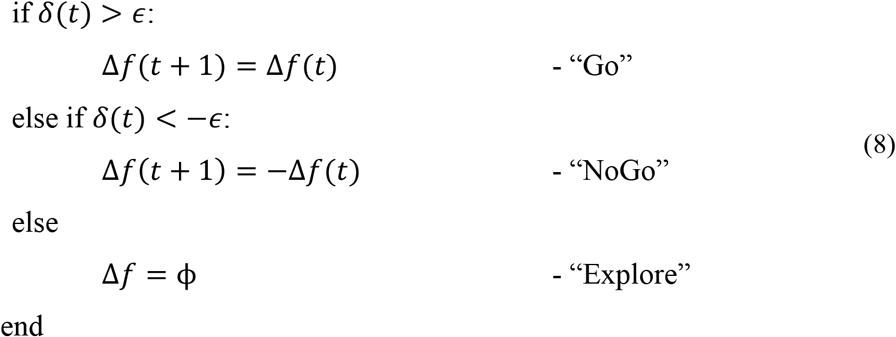

Thus, when the network receives a noisy version of the input image, the feature vector is extracted in the Central Layer of the network. The feature is modified iteratively using the above algorithm until the corresponding familiarity value reaches the maximum. Once the familiarity value attains 1 (or the local maximum), then the latest modified feature is given to the decoder, and the output image for the proposed model is produced.

#### 2.1.9 Performance Comparison

The performance of the above three networks is compared on the pattern completion task for the same set of images.

For the standard convolutional autoencoder, the output image for the given input image is taken while no hill-climbing dynamics is applied over the encoded feature vector.

The recurrent autoencoder (Autoencoder with the outer loop, where the output for the current iteration is given as input for the next iteration) forms a loop, and therefore the network acts as an attractor. In this case too, there is no additional dynamics applied to the encoded feature vector, which is the output of the Central Layer. So, the output settles at a particular image output over multiple iterations for a given input image. This settled image pattern is taken as the final output of the network.

The Attractor-based Autoencoder model retrieves the output after applying the familiarity dynamics to the Central Layer output using the familiarity value. The familiarity value, V, is extracted from the single node using the feature vector from the Central Layer (Figure 3). Figure 6 shows the familiarity value concerning the noise percentage for images of zero at different noise levels. Here the actual familiarity value is derived using equation (1). The predicted familiarity value is the output of the single value node.

**Figure 6.**
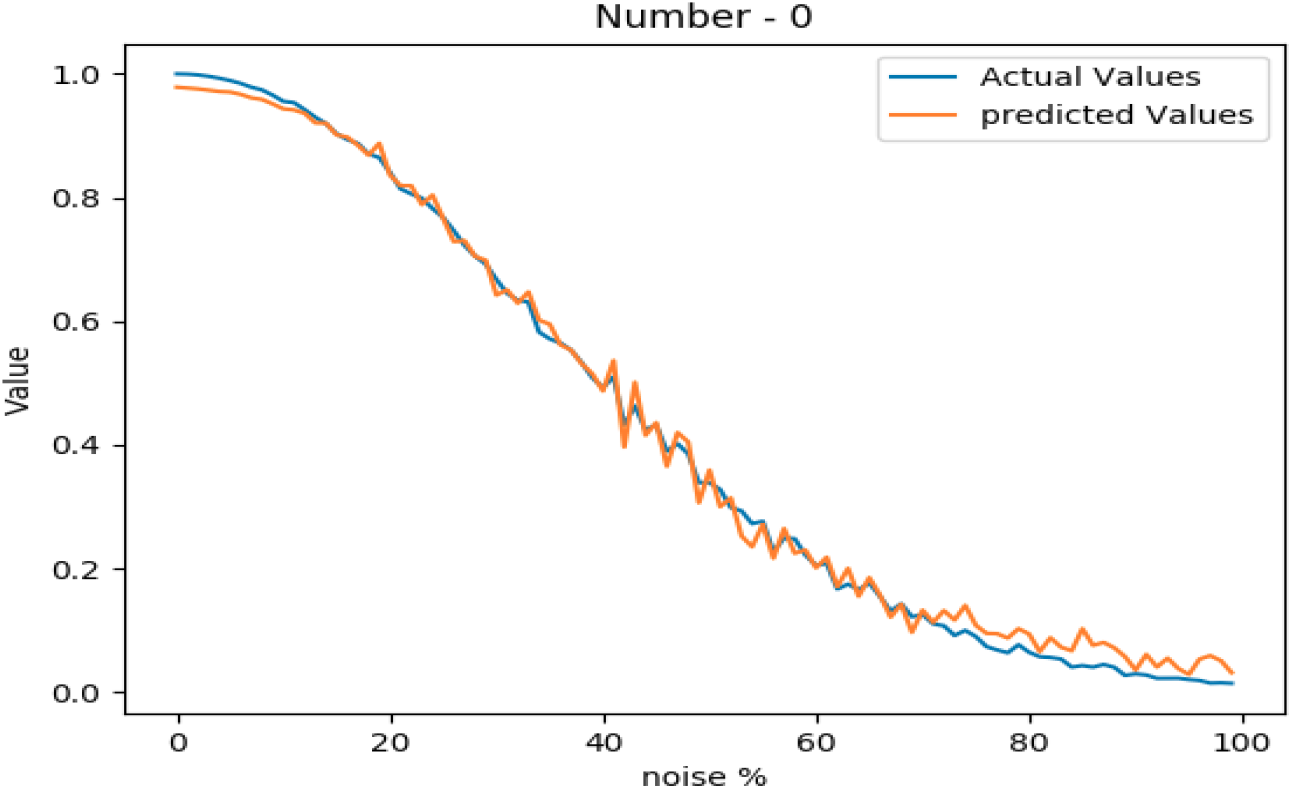
Comparison between the actual familiarity value and the network predicted value at different noise levels

#### 2.1.10 Results

The performance on the pattern-completion task is compared here for the above three models. Figure 7 shows the output comparison at different noise levels. The first row has the input images of “3” at five different noise levels. The second, third, and final rows show the outputs of the standard (SCA), recurrent (RCA), and the proposed Attractor-based convolutional autoencoder Model (ACA), respectively. It clearly shows that the network with the GEN technique outperforms both the other methods in reconstructing the proper images from the noisy images.

**Figure 7.**
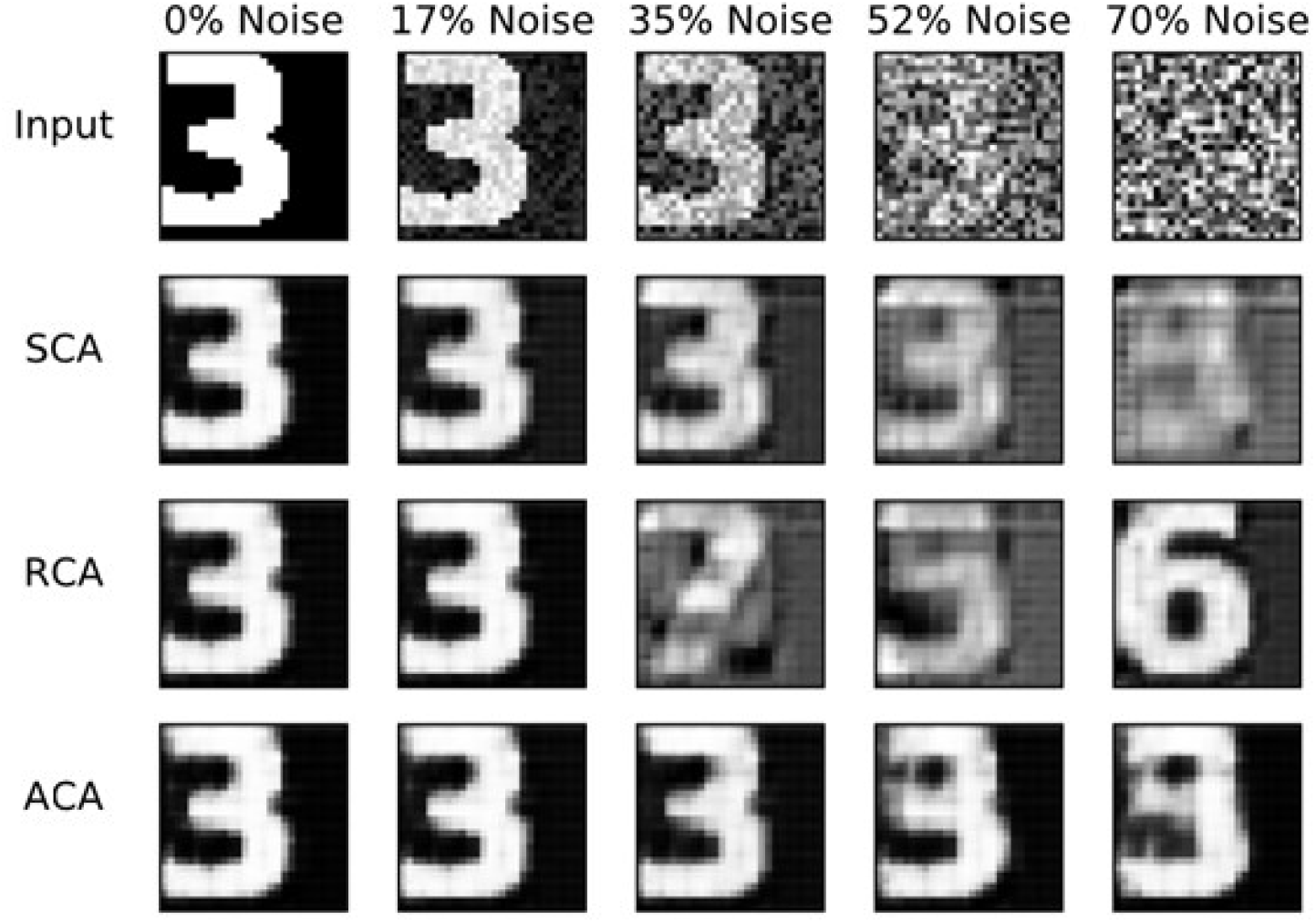
Image reconstruction comparison for Image three at different noise levels

Figure 8 compares the noise reduction/removal capability (RMS error) among the three models. The RMS error is estimated using the equation (9).

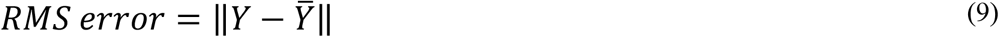

where *Y* is the expected noiseless Image and 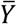 is the network output. Even at higher noise levels, the ACA model retrieves better noiseless images. It explains the need for an inner loop that estimates and uses the familiarity function.

**Figure 8.**
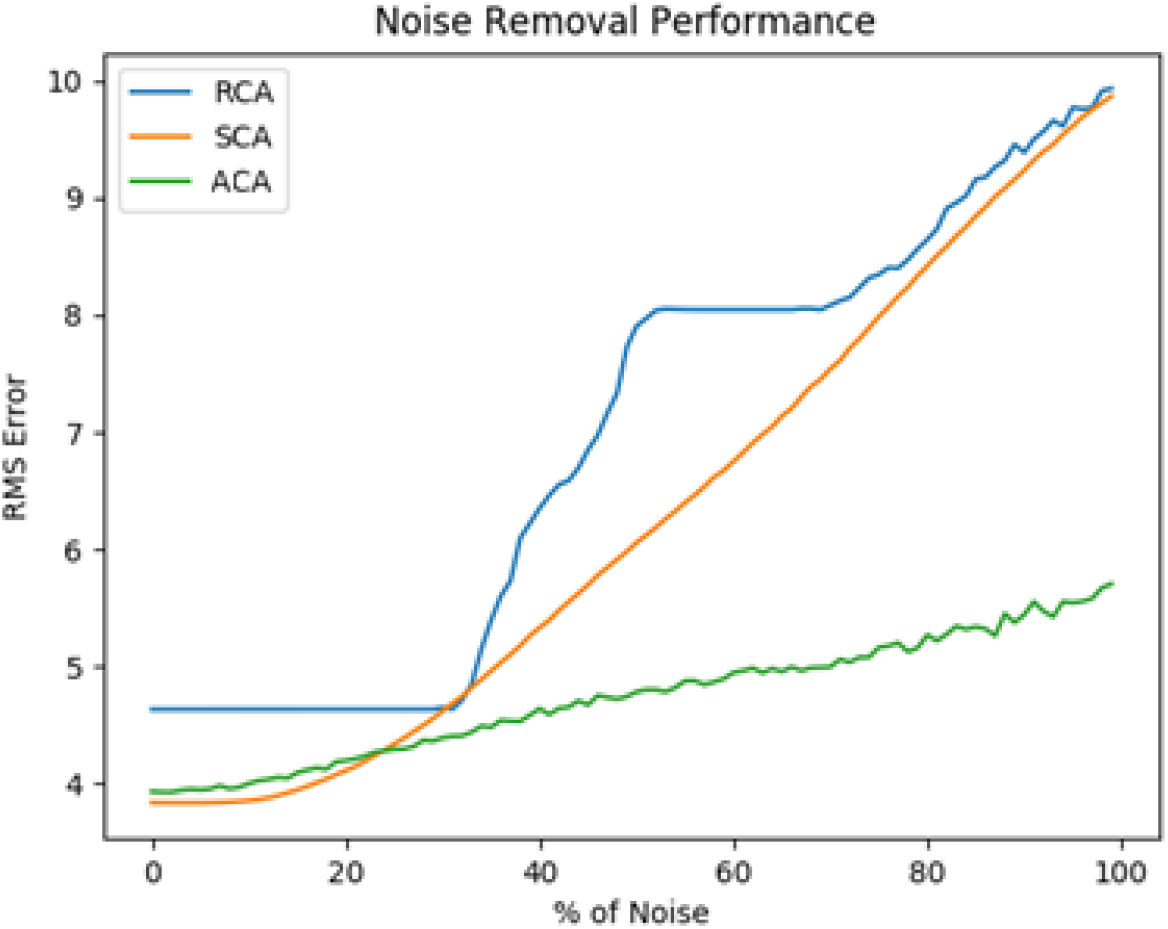
Comparison of Reconstruction error between SCA, RCA, and ACA at different noise levels

### 2.2 Hetero-Associative Memory Model

The hetero-associative memory model is demonstrated using a multimodal autoencoder network (Ngiam et al. 2011). The network used here is an extension of the value-based convolutional autoencoder. The hetero-associative memory behavior is instantiated in the image-word association task, which can be compared to the behavior of AD patients in the picture-naming task.

#### 2.2.1 Multimodal autoencoder network

The multimodal autoencoder network has two components, the Image Autoencoder and the Word autoencoder. Both the components here are joined at a Central Layer (Figure 9). The Image Autoencoder and the Word autoencoder take images and words as inputs respectively. A similar network configuration was used in another model called the *CorrNet* to produce common/joint representations (Chandar et al. 2016). The Image encoder uses two convolution layers with max-pooling followed by two fully connected layers. The Word encoder uses three fully connected layers. The outputs of the image encoder and word encoder are combined to make a common/joint representation. From this common representation layer, a single neuron is connected via a sigmoidal layer with 32 neurons, which outputs a scalar value representing the familiarity level of the input. The image decoder with two fully connected layers followed by two deconvolutional layers uses the joint representation to generate the image output. Similarly, the word decoder with three fully connected layers uses the joint representation to generate the Word output.

**Figure 9.**
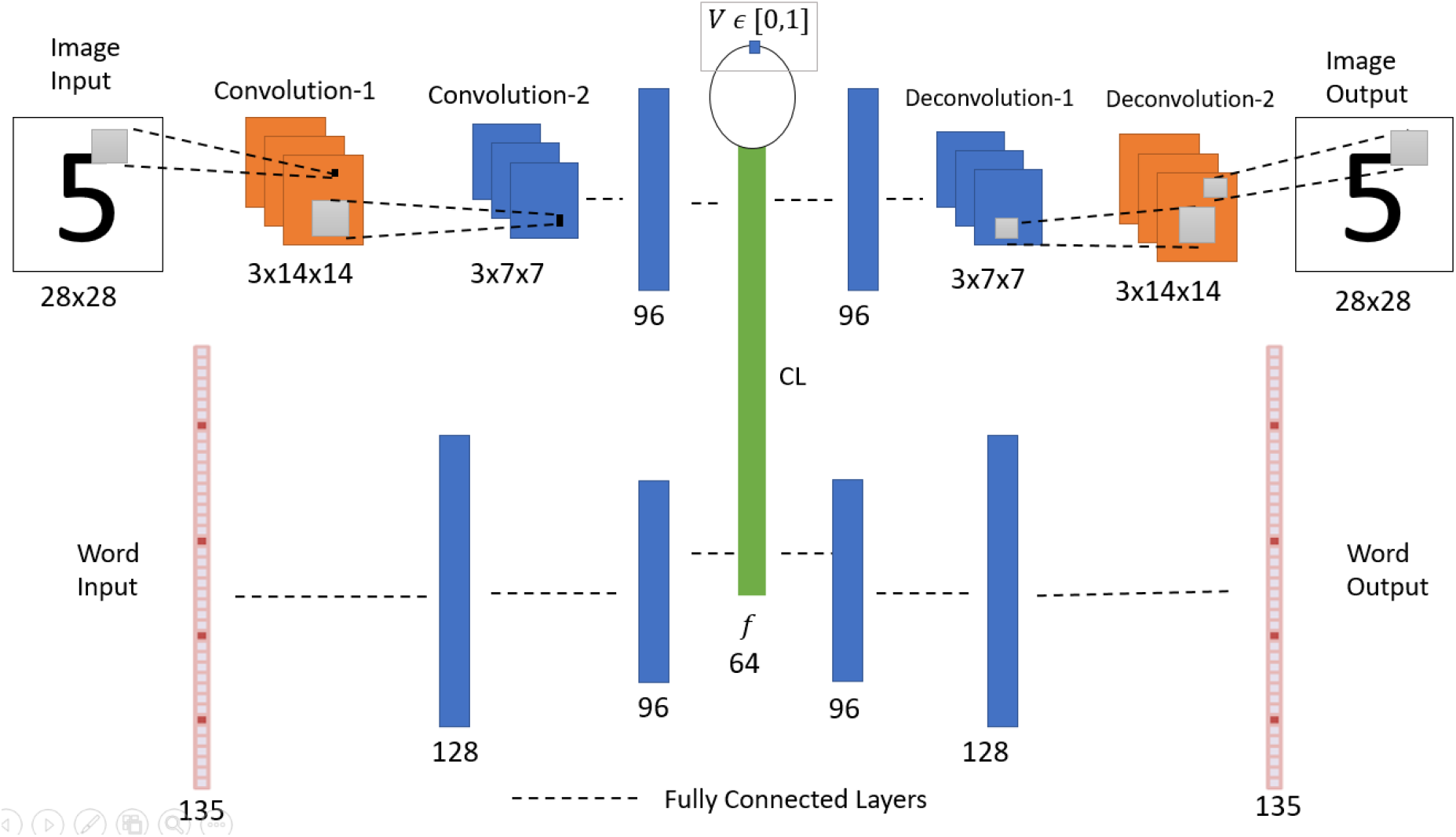
Network Architecture of Multimodal autoencoder with an associated value function. CL-Central Layer

The robustness of associative memory behavior is tested by resetting the neurons at the Central Layer at different percentage levels for a given input image and generating the image and word outputs.

#### 2.2.2 Image Encoder

Similar to the previous convolutional autoencoder network, the image encoder uses two convolution layers with the same number of filters of the same size. The second convolutional layer output is connected to a fully connected layer with 96 neurons. All the hidden layers use the leaky-ReLU activation function (Maas, Hannun, and Ng 2013).

#### 2.2.3 Word Encoder

The Word input is processed by the encoder with fully connected layers. The encoder takes a vector of size 135 as input. This vector represents five characters each by a 27 sized vector, thus 5×27=135. A detailed explanation of this vector is given in the dataset section. The input layer is connected to the first layer with 128 neurons. The first layer is connected to the second layer with 96 neurons.

#### 2.2.4 Joint representation

The outputs of both the image encoder and the word encoder are connected to a layer with 64 sigmoidal neurons. This generates a common feature. The common feature is estimated using the following equation (10).

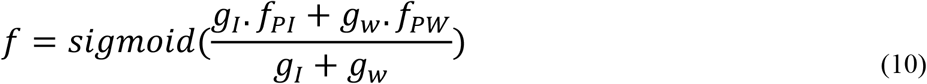

where *f_PI_, f_PW_* denotes the pre-feature vector (before applying activation function) from the Image encoder and the Word encoder, respectively; *g_I_* and *g_w_* are binary values representing the availability of image and word inputs, respectively, where 1 denotes the availability of a particular input, and 0 denotes the non-availability of input.

#### 2.2.5 Familiarity Value function

The Central Layer is connected to a single sigmoidal neuron via a single hidden layer with 32 sigmoidal neurons. The single output sigmoidal neuron estimates a scalar value denoting the familiarity level of the input image-word combination.

#### 2.2.6 Image Decoder

The Image decoder uses the same architecture as in the standard convolutional autoencoder network.

#### 2.2.7 Word Decoder

The Word decoder takes the common feature as input and processes it through 3 fully connected layers that have 96, 128, and 135 neurons, respectively. The first two layers use the leaky-ReLU activation function. The output layer uses 5 SoftMax functions, out of 27 neurons each. Here each SoftMax function specifies one character.

#### 2.2.8 Dataset

The dataset used here is a combination of Images, Words, familiarity values, and association indices. The images are used from the dataset as in the autoencoder network.

#### 2.2.9 Words

The Word input is represented using five numbers of 27-dimensional one-hot vector representations, which together form a vector of size 135 (= 27 × 5). A single character is represented by a 27-dimensional vector. Among the 27 dimensions, the first 26 dimensions represent alphabets (a-z), and the last (27th) dimension represents the special character - empty space. For example, for the character ‘e,’ the 5th element in the vector is set to 1, and the rest of the elements are set to 0. The number of characters is chosen to be five, considering the maximum number of characters in the words for numbers from zero to nine.

The Word input data is generated for the number-names (*zero, one, two*,…., *nine*) and the number-type-names (*even*, *odd*) (Table 1).

**Table 1.** Word Inputs

| Number-Names |  |
| --- | --- |
| Zero | One |
| Two | Three |
| Four | Five |
| Six | Seven |
| Eight | Nine |
| **Number-type names** |  |
| Even | Odd |

The noisy words are generated using equation (11). This way,12,000 noisy words are generated and used.

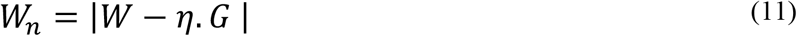

Where *W*, and *W_n_* denotes the proper and noisy Words, respectively. *G* is a noise vector with dimension 135, and each element of which is sampled from the standard uniform distribution *U*(0,1).

#### 2.2.10 Familiarity Value

The familiarity value for both the Image and Word is calculated using the Gaussian formula as in Equations (12) and (13).

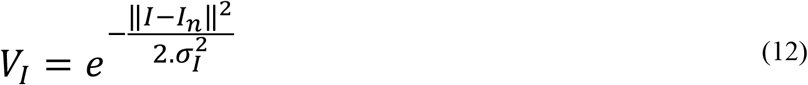

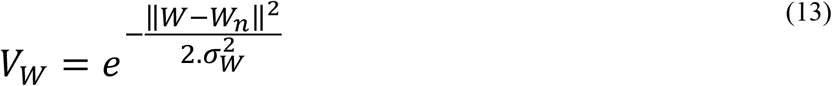

where *I, W, I_n_, W_n_* denotes the noiseless Image, noiseless Word, noisy Image, and noisy Word, respectively. Here *σ_I_* is set to 50 and *σ_W_* is set to 8. Thus, a noise-free image/word has a familiarity of 1; when there is noise, it will have a value between 0 and 1 depending on the level of noise.

When both the image and word inputs are presented to the network, the combined Familiarity is calculated by multiplying the familiarity value of the Image and the Word. This familiarity value is used by the network to reach the nearest best feature with maximum value by the Go-Explore-NoGo technique.

#### 2.2.11 Association Index (*γ*)

The association index, ***γ***, is a scalar that specifies the relation between the Image and the Word. For various combinations of the above images and words, the association index is generated. The rules followed for generating the association index are listed below.

- If a word denotes the same number-name or number-type-name for the given Image, then *γ* is set to +1.
- If a word does not match with either the number-name or number-type-name for the given Image, then *γ* is set to −1.

For example, ‘0’ (Image) and ‘zero’ (word) have an association index of +1, similarly ‘0’, and ‘even’ also get the association index of+1. But, ‘0’ and ‘odd’ have an association index of −1. At a particular instant, at least one among the Image or word data should be present. When one modality among Image and Word is absent, the association value is set to 0.

Along with all the above input data, two more binary values *g_I_* and *g_W_* are also used, which represent the presence of the image input and the Word input, respectively.

#### 2.2.12 Training

For training the network, a combination of Image, Word, familiarity value, and the association index are used. The three input-output combinations used for training the network are shown in Table 2.

**Table 2.** Input-Output Combinations for the Multimodal Autoencoder

| Input | Output |
| --- | --- |
| Image, Word | Image, Word, Combined Familiarity Value, and Correlation-coefficient |
| Image | Image, Image Familiarity Value |
| Word | Word, Word Familiarity Value |

The multimodal network is trained to produce the same given inputs as outputs irrespective of the noise in the input.

Different Image-Word combinations are used for training the network. The various combinations are as follows:

1. Noiseless Image, noiseless Word
2. Noisy Image, noiseless Word
3. Noiseless Image, noisy Word

Here’ noisy image and noisy word’ combination is not used for training.

Among the above three combinations in the training dataset, each of the above combinations has 100, 1,00,000, and 12,000 data, respectively. From the above combinations, each training batch contains 100, 1000, and 120 data, respectively. Thus, each batch contains 1220 data. Here all the data points in the first combination are used in all the batches to match the count, which is high for the other data combinations. The network is trained for 10,000 epochs.

The network is trained to minimize the cost function 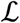 (Equation (19)). A combination of 5 cost functions 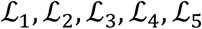 are used for training. 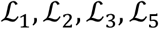 are image reconstruction error, word reconstruction error, the correlation coefficient of Image pre-feature and Word pre-feature, and value reconstruction error, respectively. 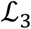 is used to form the relationship between the features of the Image and the Word. 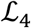 is used to make the Image and Word feature representations closer.

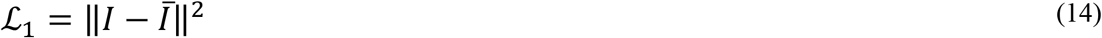

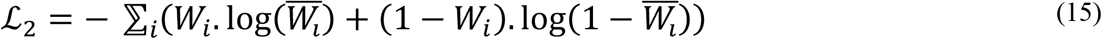

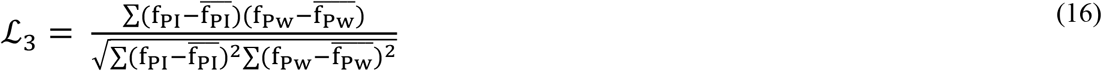

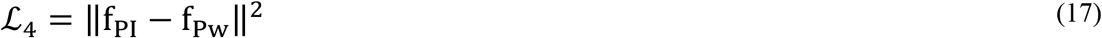

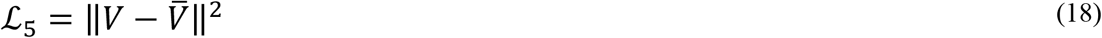

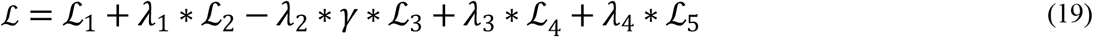

where,

I – input image
W – input word
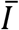 - predicted Image
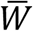 - predicted Word
f_PI_ - the pre-feature vector for Image
f_Pw_ - the pre-feature vector for the Word
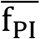 - the mean pre-feature vector for Image
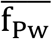 - the mean pre-feature vector for the Word
*λ*_1_, *λ*_2_, *λ*_3_, *λ*_4_ - tradeoff parameters
*γ* – association index
V – desired familiarity value
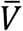 – predicted familiarity value The network parameters are updated using Adam optimizer (Kingma and Ba 2015). After training, the results are generated with various combinations of inputs.

#### 2.2.13 Results

The results are generated by giving only one input (either Image or word) at a time. Figure 10 visualizes the common feature vector (of size 64) in 2D space (using the best two components of Principal Component Analysis (PCA)). The Word inputs are given to the network while keeping the image inputs blank (zero values). 0-9 specifies words’ zero,’ ‘one,’…, ‘nine’ respectively, where ‘10’ corresponds to the word ‘even,’ and ‘11’ corresponds to the word ‘odd.’ Note that the words corresponding to the odd type and even type form separate clusters.

**Figure 10.**
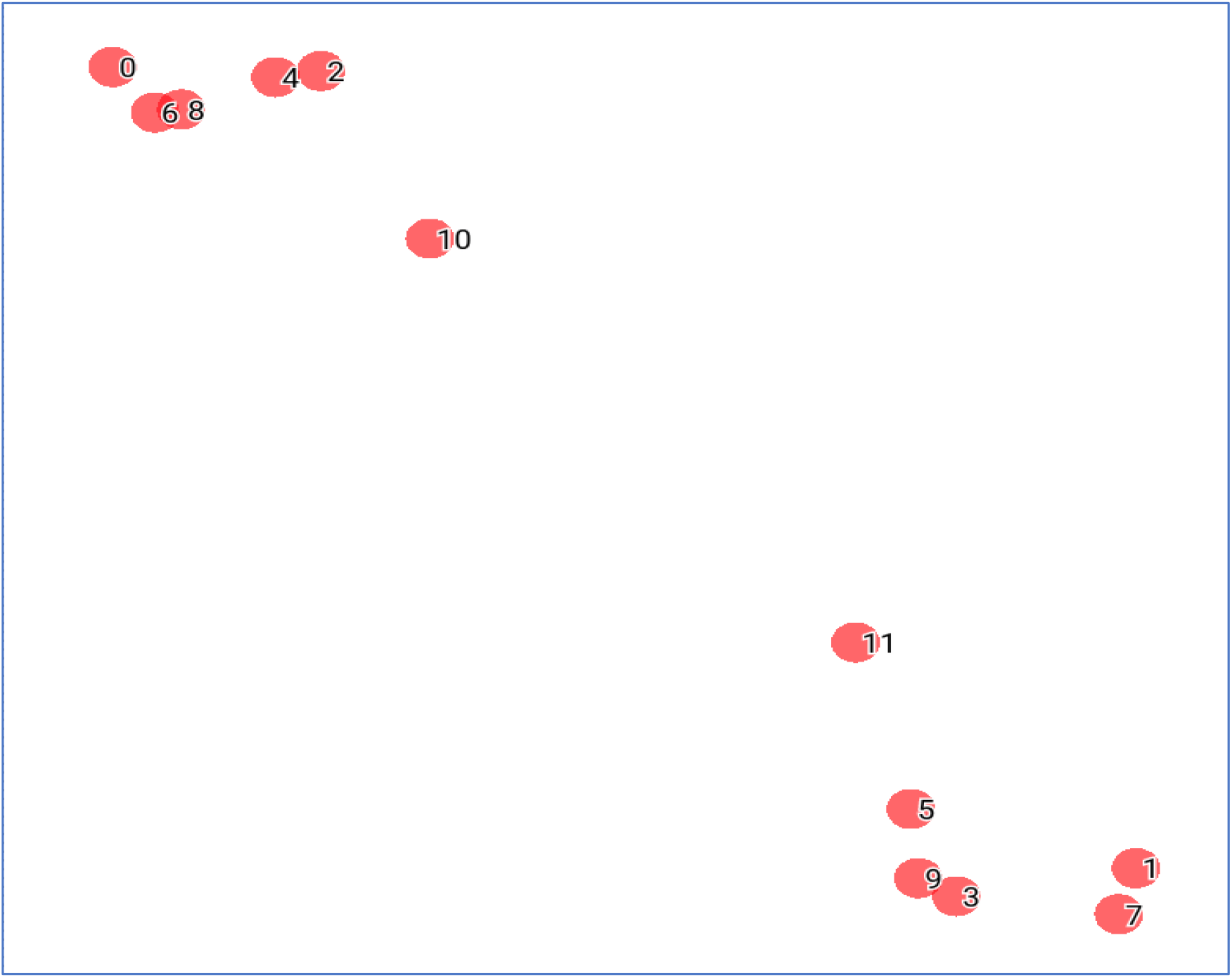
Vector representation of word features in 2D space

The results below (Figure 11–Figure 15) are generated by giving image input alone while keeping the Word input to be empty (zero values).

**Figure 11.**
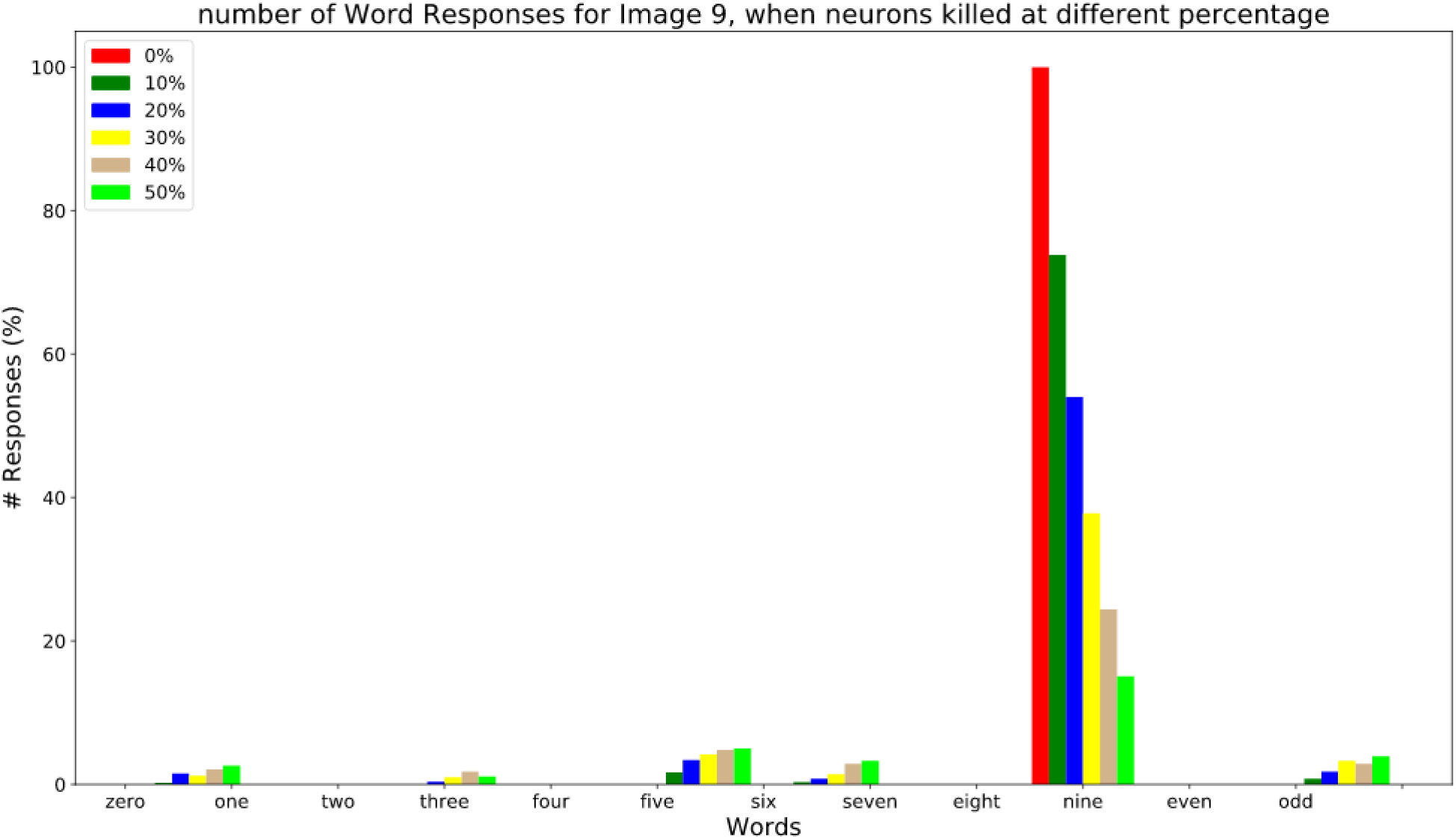
the response count for all the number-names and the number-type-names while resetting different percent of neurons (0%, 10%, 20%, 30%, 40%, 50%,60%) for the image input of number 9.

**Figure 12.**
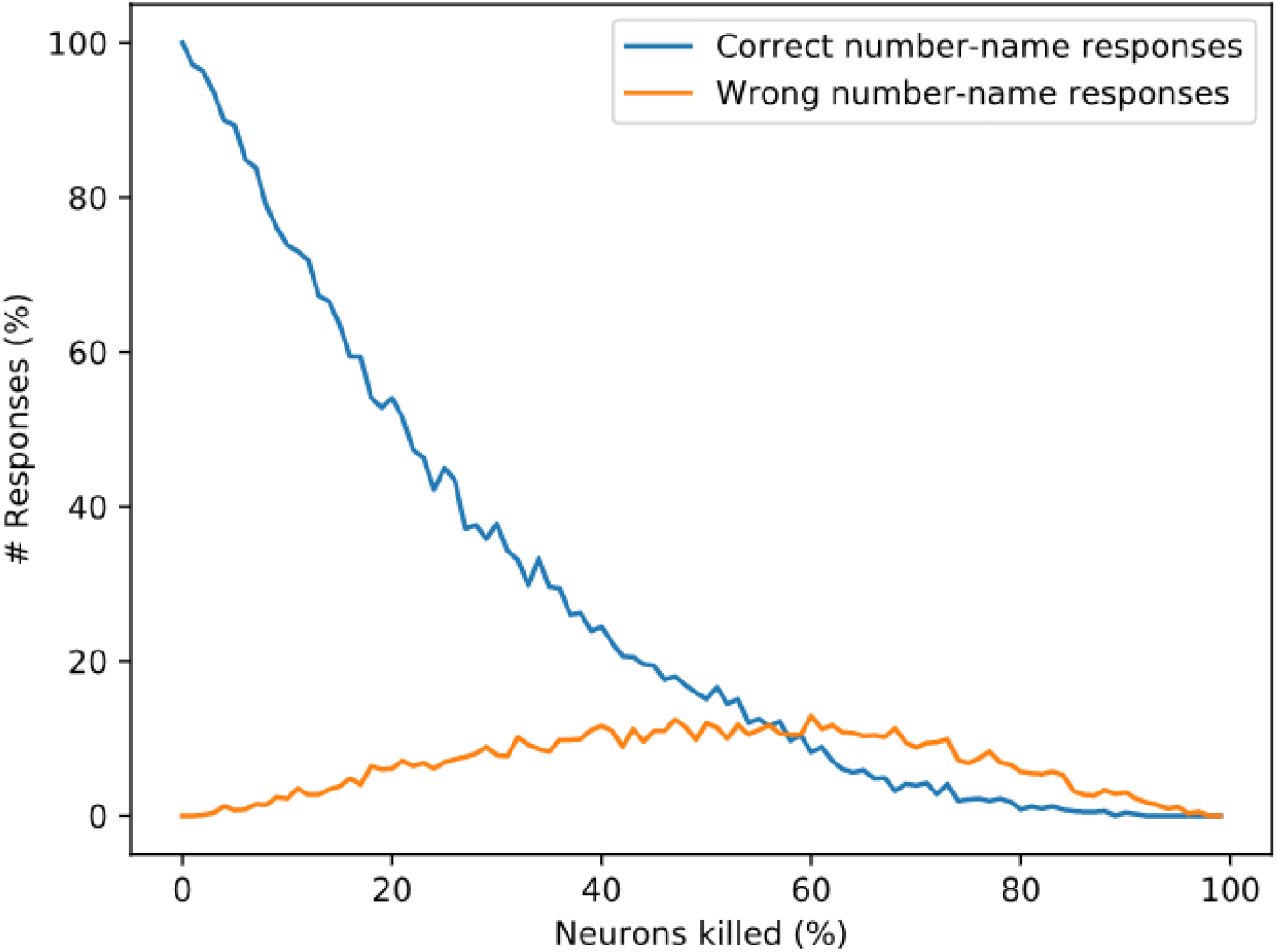
response percentage comparison of correct vs. wrong number-names

**Figure 13.**
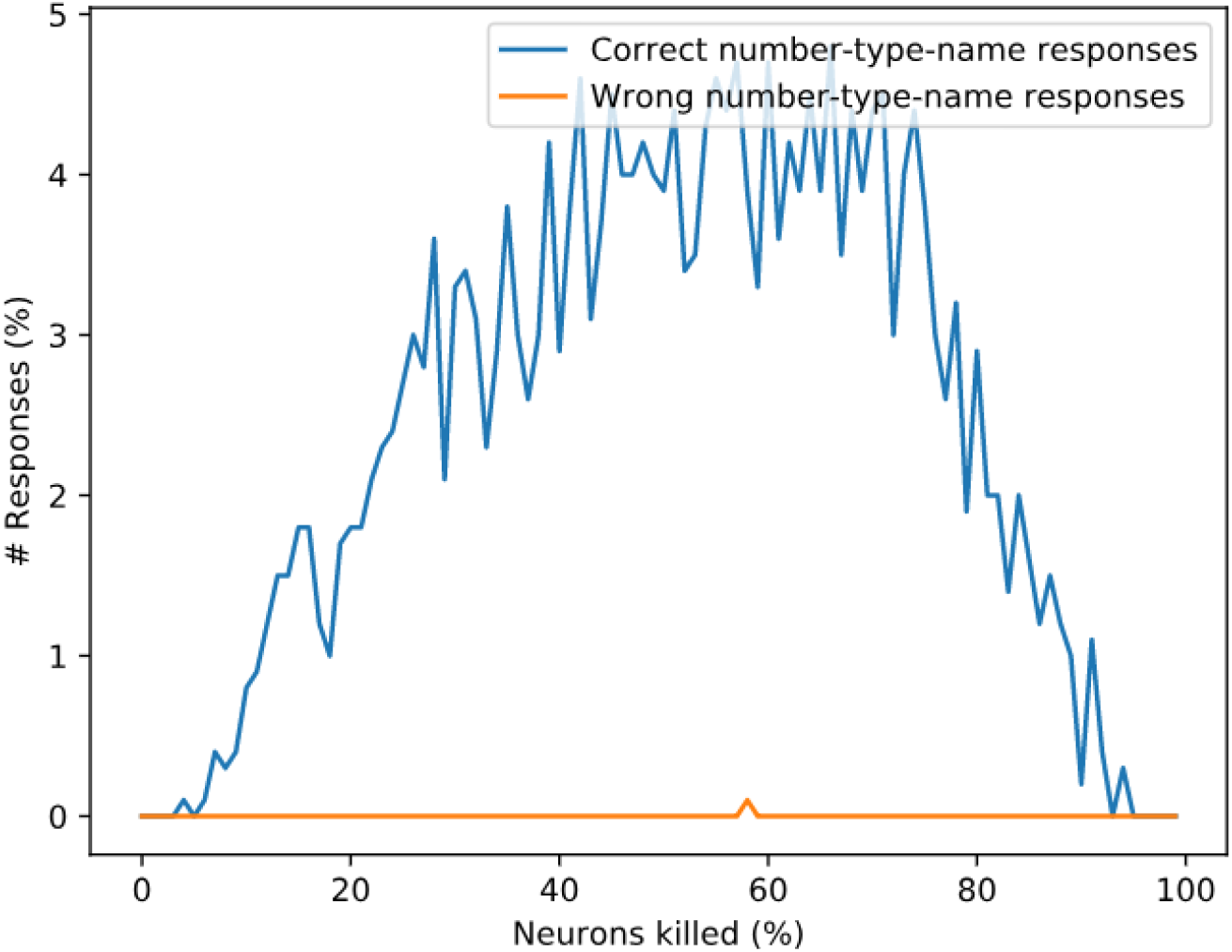
Comparison of percentage of correct number-type-name vs. wrong number-type-name responses for image input of ‘9’

**Figure 14.**
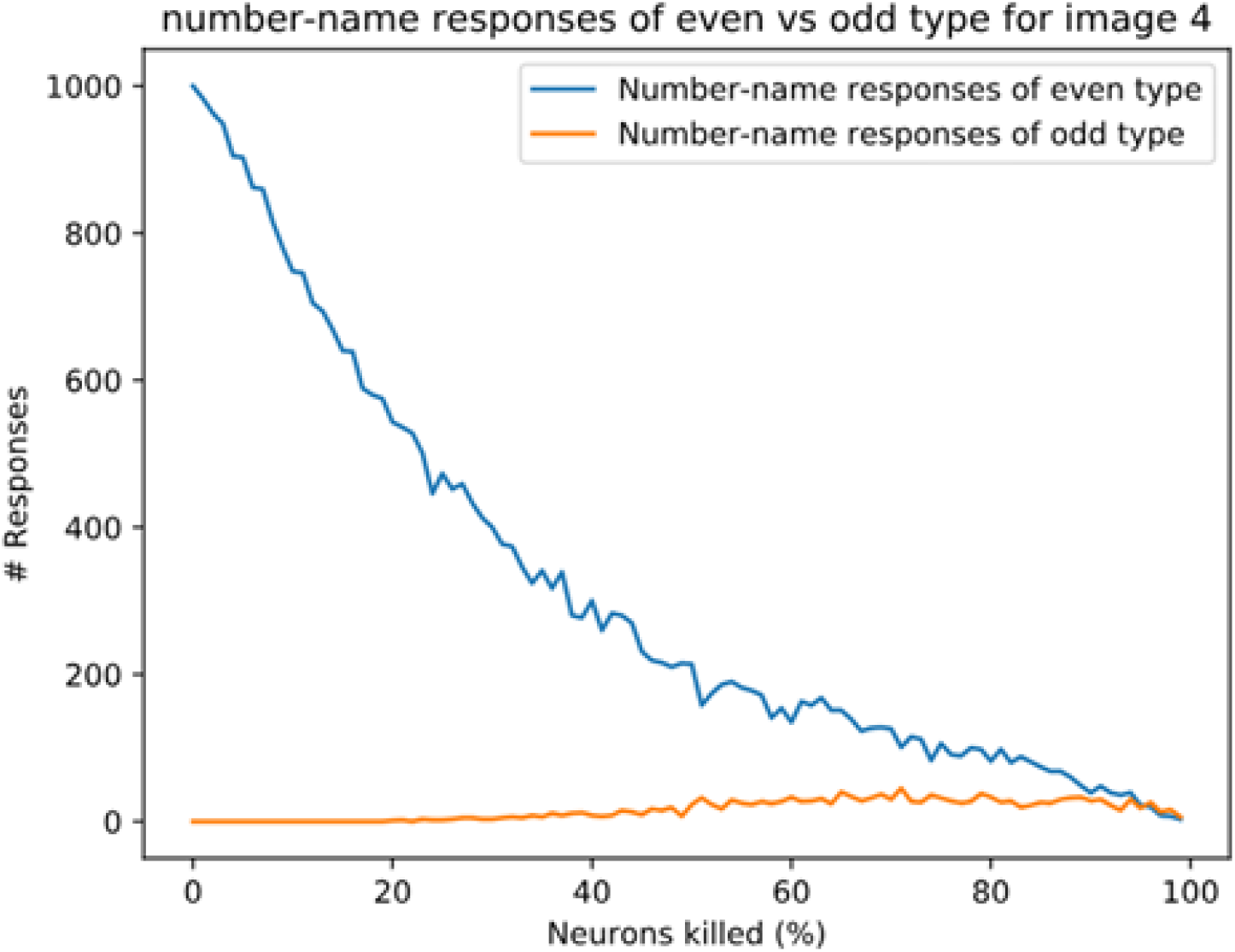
Sum of the count of even number-name responses vs. odd number-name responses for image input 4

**Figure 15.**
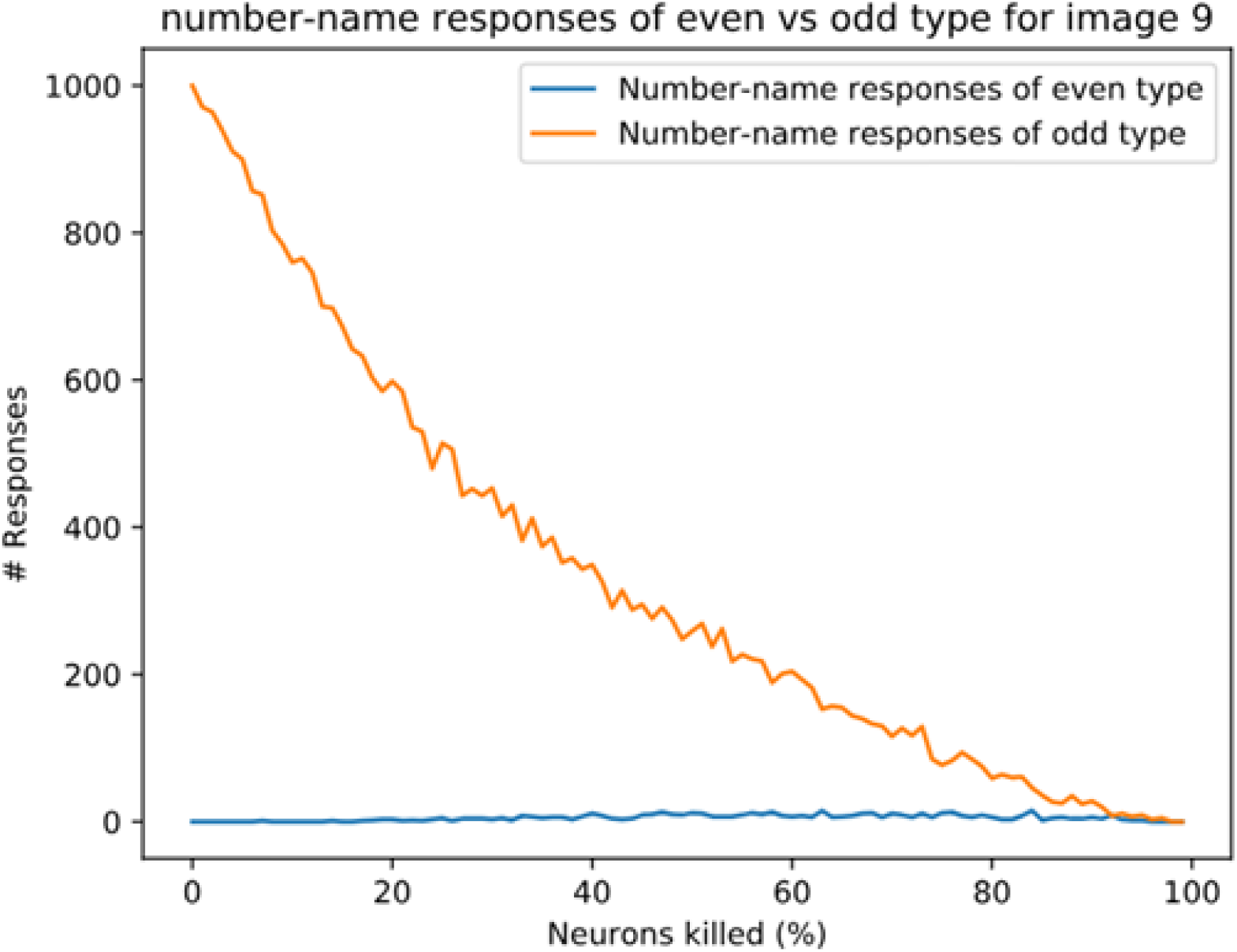
Sum of the count of even number-name responses vs. odd number-name responses for image input ‘9’.

#### 2.2.14 Simulating the Behavior of Alzheimer’s Disease (AD) patients on Picture-naming Task

AD is characterized by the loss of cells in the Entorhinal cortex, a cortical area that serves as the gateway to the hippocampus (Gómez-Isla et al. 1996; Bobinski et al. 1998). Since the Central Layer, and the associated Familiarity computation network represent the Hippocampus in the proposed model, AD pathology is simulated by randomly killing/resetting a percentage of the neurons in the Central Layer. Then Go-Explore-NoGo policy is applied over this modified feature to reach the feature with maximum familiarity value. Note that the killed/reset neurons do not participate in the computation. The graphs below are generated by counting each word output’s responses at the Word decoder out of 1000 times.

We consider three kinds of responses for a given image input while resetting neurons at the Central Layer. They are number-name responses (zero, one,. nine), number-type-name-responses (even, odd), and non-name responses (anything other than number-names and number-type-names).

The response percentage of all the number-names and number-type-names for the image input ‘ 9’ is shown in Figure 11. In order to simulate AD pathology at different levels of degeneration, in the Central Layer, different percent of neurons (0%, 10%, 20%, 30%, 40%, and 50%) are reset, and the average response count is calculated. From this, we can observe that as the percent of neurons being reset increases, the correct responses decrease. When the image input ‘9’ is presented, and no Central Layer neuron is reset, the network produces the word “nine” all the time as expected. When 10-30% of neurons are reset, it produces the word “nine” most of the time. Among the wrong responses, most of them are either number-name responses or number-type-name responses of the same type/category (in this case: one, three…, nine, and odd). In other words, the responses of number-names (one, three, five, seven, and nine) and number-type-name (odd) of the same group are high compared to the number-names (zero, two, four, six, and eight) and number-type-name (even) of a different group. This can be related to the semantic error. When 40-50% neurons are reset, the sum of all the number-name and number-type-name responses falls below 30%, and the non-name response count is higher, which is similar to No-response.

Figure 12 shows the count of correct number-name response (‘nine’) and wrong number-name responses (all the number-names except ‘nine’) while resetting different percent of neurons for the input of image “9”. This doesn’t include the number-type name responses such as ‘odd’ and ‘even.’

Figure 13 shows the count of the correct number-type-name response (‘odd’) and wrong number-type-name response (‘even’) while resetting different percent of neurons for the input of image “9”. The number-name responses such as ‘zero,’ ‘one,’ etc., are not counted here.

From Figure 12 and Figure 13, We can observe that, as the percentage of neurons being reset increases, the response count of correct number-names (nine) reduces gradually, whereas the response count of the wrong number-names and the correct number-type-name (odd) increase gradually for some time and decrease after that. Here among the wrong number-name responses, most of them are of the same type but different number-name responses (one, three, five, seven), which can be related to the semantic error.

Figure 14 and Figure 15 show the count of even number-name responses vs. odd number-name responses (excluding the number-type-names) for the image input ‘4’ and ‘9’, respectively. It can be observed that, for a given image input, the chance of producing number-name word response of the wrong category is very small. This also explains the logic behind the occurrence of semantic errors.

## 3. Discussion

We present a deep network-based model of the associative memory functions of the hippocampus. The cortico-hippocampal connections are abstracted out into two structural modules of the proposed model. In the first module, the bidirectional cortico-hippocampal projections are modeled as an autoencoder network. In this second module, the loop of connections from EC into the hippocampal complex and back to EC is modeled as hill-climbing dynamics over a *familiarity* function.

In the first part of the study, the model is used to simulate auto-associative memory functions using pattern completion task under normal conditions. The pattern-completion task is modeled using a convolutional autoencoder with an associated familiarity function. The autoencoder’s encoder and decoder are related to the feedforward and feedback projections between the sensory cortices and the hippocampal formation. There are many conventional denoising autoencoders proposed to solve this problem (Tian et al. 2019). These models use a supervised learning approach, where the noisy patterns are mapped to noiseless patterns during training. This kind of mapping does not fit the actual scenario, where the brain is not always presented with noisy and noise-free versions of the same pattern. The present study maps the noisy patterns to the same noisy version itself. The model learns to construct a noise-free version on exposure to a large sample of noisy patterns.

### 3.1 Familiarity and the Hippocampus

The concept of familiarity figures invariably in most discussions of the memory functions of the hippocampus. Studies on human memory that draw from cognitive, neuropsychological, and neuroimaging methodologies suggest that human memory is composed of two processes of memory: *recollection* and *familiarity* (Henson et al. 1999; Andrew P Yonelinas et al. 2005; Andrew P Yonelinas 2001; Droege 2017). Sometimes when we meet a person, we may simply have the sense that the person is familiar but not remember the person’s name or when and where we have first met that person. This sense of having-met-before refers to familiarity, while the ability to recall the various properties that constitute that object refers to recollection. Four empirical features were thought to dissociate familiarity from recollection. Familiarity and recollection differ in terms of 1) speed of processing, 2) Receiver Operating Characteristics, 3) electrophysiological signatures, and 4) neuroanatomical substrates (Andrew P. Yonelinas 2002).

Several proposals were made regarding neuroanatomical substrates of familiarity and recollection. Based on the memory performance of patients with medial temporal lobe damage, some researchers suggested that while the hippocampal region is necessary for recollection, the surrounding cortical structures like the parahippocampal gyrus are essential for familiarity (Huimin, Aggleton, and Brown 1999; Eichenbaum, Otto, and Cohen 1994). Another proposal links recollection with medial temporal lobe structures and familiarity with existing memory representations in the neocortex (J. M. Mandler and DeForest 1979; G. Mandler 1980; 1991; Graf, Squire, and Mandler 1984; Graf and Mandler 1984). Other proposals suggest that the hippocampus is important for both the familiarity and recollection processes (Wais et al. 2006; Malmberg, Zeelenberg, and Shiffrin 2004). Manns et al. (2003) showed that the lesion limited to the hippocampus impaired recollection and familiarity in the remember/know procedure (Manns et al. 2003). Wixted et al. (2010) show that familiarity is supported by the hippocampus when the memories are strong (Wixted and Squire 2010). They also argue that the hippocampus and the adjacent regions do not exclusively support only one process (Wixted and Squire 2010). Although several authors accept the existence of dual processes – recollection and familiarity - there is no consensus on the neural substrates of recollection and familiarity. We now present a neurobiological interpretation of the computation of familiarity in the hippocampal circuitry. Mesencephalic dopaminergic signals have a major role to play in the proposed theory.

### 3.2 Dopamine, Memory, and Hippocampus

The importance of dopamine in memory encoding and recall has been observed in many studies. Although dopamine signaling, in the context of the BG, is often associated with motor function, there is extensive evidence linking dopamine to cognition and memory functions (Goldman-Rakic 1997; Kulisevsky 2000; Koch et al. 2014; Martorana and Koch 2014).

There is a considerable body of neurobiological literature that links dopaminergic signaling with reward processing (Wise and Rompre 1989). Using classical conditioning experiments, Schultz et al. (1997) took a further step and demonstrated strong analogies between dopaminergic activity in Ventral Tegmental Area (VTA) and an informational signal known as temporal difference (TD) error in Reinforcement Learning (W. Schultz, Dayan, and Montague 1997). This connection had inspired extensive computational modeling effort that sought to connect dopaminergic signaling with the function of the basal ganglia (BG), an important subcortical circuit linked to dopamine signaling. Dopamine projections to the striatum are thought to be responsible for the computations of value that occur in the ventral striatum (Chakravarthy and Surampudi 2010). Models of the BG constructed on the foundations of Reinforcement Learning, linking dopamine signals with reward prediction error, or small variations of such a reward-related quantity, were able to explain a wide variety of motor and cognitive functions of BG (W. Schultz, Dayan, and Montague 1997; Sukumar, Rengaswamy, and Chakravarthy 2012; Joseph, Gangadhar, and Srinivasa Chakravarthy 2010; Sridharan, Prashanth, and Chakravarthy 2006).

### 3.3 The role of dopamine in the memory functions of the hippocampus

Packard and White (1989) demonstrated memory enhancement on application of dopamine agonists (Packard and White 1989). There is strong evidence that links age-related reduction in dopamine to age-related cognitive and memory decline (Bäckman et al. 2006). Dopamine agonists, like Bromocriptine, enhanced memory performance in the elderly (Morcom et al. 2010). Even in patients suffering from Parkinson’s disease (PD), a disease thought to be predominantly motor disease, cognitive impairment is linked to dopamine-deficiency (Lebedev et al. 2014). This link is further strengthened by the application of L-Dopa, a dopamine precursor, which was found to ameliorate cognitive deficits in PD patients (Kelly et al. 2009). Similar results were obtained from animal studies also. Dopamine transporter heterozygous knockout mice showed deficits in pattern completion under partial cue conditions (Li et al. 2010).

It is possible to find neuroanatomical evidence within the hippocampal circuitry in order to support the aforementioned studies that link memory deficits with dopamine. Although there was an early view that dopamine does not modulate hippocampal neural activity, subsequently, evidence was gathered for the existence of mesencephalic dopamine projections in rat hippocampus (Gasbarri et al. 1994; Penfield and Milner 1958b). mRNA for D1- and D2-receptors were found in hippocampal neurons (Mansour et al. 1992). Dopamine modulates neurotransmission in CA1 (Hsu 1996) and CA3 (Hamilton et al. 2010) regions of the hippocampus.

Lisman and Grace (2005) presented an extensive review of experimental literature to establish not only the existence of dopaminergic projections to the hippocampus, but to show that dopaminergic neurons that project to the hippocampus fire in response to novel stimuli (Lisman and Grace 2005). Electrophysiological recordings showed that dopamine cells in VTA, projecting to the hippocampus, increased their firing rate in response to novel stimuli; the activity of these neurons reduced with increasing familiarity (Ljungberg, Apicella, and Schultz 1992; Steinfels et al. 1983). Neuroimaging studies based on PET (Tulving et al. 1992), and fMRI (Strange and Dolan 2001; Yamaguchi et al. 2004), showed increased hippocampal activity due to novel stimuli. The hippocampal activity also interfered with the rabbit’s ability to orient to the novel stimulus (Honey, Watt, and Good 1998; Vinogradova 2001). These studies clearly establish the role of the hippocampus in representing the novelty of stimuli. Considering that novelty is the complementary notion to familiarity, the above body of evidence can be invoked to support our model that requires the computation of familiarity in the hippocampus. With the above background information, the central hypothesis of the proposed model may be expressed as follows: the cortico-hippocampal interactions with regard to memory operations are based on maximizing familiarity computed within the hippocampal circuit. Since the process of memorizing a pattern entails a gradual transition from novelty to familiarity, this assumption of maximizing familiarity seems to be intuitively plausible.

In the present study, the familiarity function (equation (1)) is trained by supervised learning that involves a direct comparison of the target pattern with the predicted pattern. It is also possible to train the familiarity function by Reinforcement Learning (RL) (Sutton and Barto 2018), where a close match between the target and recalled pattern results in a reward. The familiarity function then, in mathematical terms, becomes the value function. An RL-based formulation of hippocampal memory functions has an added advantage. The reward signal can be used not only to represent the level of match between the target and recalled pattern but also to represent the saliency of the pattern to the animal/subject. Several existing accounts that posit CA3 as the site of memory storage in the hippocampus, argue that the decision to store or not to store depends solely on the mismatch between the target and stored pattern (Hasselmo, Schnell, and Barkai 1995; Treves and Rolls 1992). But a memory mechanism that stores all novel stimuli encountered by the animal in its interactions with the world, irrespective of the saliency of the stimuli to the animal, would glut the animal’s memory resources. The best possible way is to store only the important stimuli by filtering in based on the salience factors such as reward, novelty, recency, and emotional involvement (McNamara et al. 2014; Singer and Frank 2009; Cheng and Frank 2008). These salience-based mechanisms permit the storage of patterns with high relevance (Santangelo 2015). It is also important to notice that the dopaminergic nuclei projected to the hippocampus support the dopamine release by novelty and/or reward-related experience, which adds meaning to the processed information (Lisman and Grace 2005; Hansen and Manahan-Vaughan 2014). These notions will be explored in our future efforts.

The possibility of using RL concepts to describe hippocampal memory operations brings up an interesting analogy between the hippocampus and another subcortical circuit – the basal ganglia (BG). The BG has been modeled by several researchers using concepts from RL (Sukumar, Rengaswamy, and Chakravarthy 2012; Sridharan, Prashanth, and Chakravarthy 2006; Joseph, Gangadhar, and Srinivasa Chakravarthy 2010). In fact, the hill-climbing algorithm for maximizing familiarity in the proposed study was earlier used in a different context (Srinivasa Chakravarthy and Balasubramani 2018). The algorithm, known as Go-Explore-Nogo (GEN) policy, was used in a model of the basal ganglia to find optimal actions by maximizing the value function. In the context of spatial navigation, there is extensive experimental and computational literature that shows how the two subcortical systems closely cooperate in navigating the animal through its spatial environment (Fox et al. 2009). In spatial navigation literature, computational modelers have applied RL tools to describe the functions of the hippocampus and the BG (Sukumar, Rengaswamy, and Chakravarthy 2012; Chavarriaga et al. 2005; Dollé et al. 2010; Trullier et al. 1997). These potential analogies will be explored more deeply in our future models of hippocampal memory operations.

### 3.4 Modeling Hetero-associative memory function in Alzheimer’s Disease (AD)

The second part of the work demonstrates hetero-associative memory using a multimodal autoencoder. In this case, the network is trained to form association between images and words at the Central Layer. Here the trained features belonging to the same category form a cluster (even and odd). AD patients’ behavioral response during the picture-naming task is reproduced by killing/resetting the neurons at the Central Layer.

The picture-naming task is a classic example of hetero-associative memory (Cuetos, Gonzalez-Nosti, and Martínez 2005; Barbarotto et al. 1998). In this task, the participants are asked to name the pictures shown on the screen. This task is used to assess the level of cognitive deterioration in AD patients. AD is a progressive neurodegenerative disorder in which neuronal loss is observed throughout the brain. The initial loss of neurons is detected in the Entorhinal cortex and hippocampus (Gómez-Isla et al. 1996; Bobinski et al. 1998). Alzheimer’s patients at different stages show different kinds of responses in the picture-naming task. The controls and early-stage AD patients predominantly produce correct responses (e.g., to the picture of a lion, they respond with the word ‘lion’). In the mild to moderate stage, they make some semantic errors (e.g., responding with words that are similar or closer to the actual word semantically – like *tiger* in place of *lion* - or the superordinate words - like *animal* instead of *lion*). In the severe stage, they predominantly make Semantic Errors or No Response (*I don’t know*) (Cuetos, Gonzalez-Nosti, and Martínez 2005; Barbarotto et al. 1998).

In the proposed model, the severity of AD is related to the percentage of neurons killed at the Central Layer, which represents the Entorhinal cortex. The output of the network matches the behavioral response of AD patients at different levels of severity. The intact network produces the correct number-name responses for the given image input, which matches the controls and AD patients at an early stage. The network with a lower percentage of neurons killed demonstrates semantic error responses (correct number-type name response or wrong number-name of the same type), which matches the mild-moderate stage of AD patients. A high percentage of neuronal loss shows semantic errors and no response (non-name response) that matches the response of the severe stage of AD patients (Cuetos, Gonzalez-Nosti, and Martínez 2005; Barbarotto et al. 1998).

